# Bayesian inference of phylogenetic distances: revisiting the eigenvalue approach

**DOI:** 10.1101/2024.03.27.586929

**Authors:** Matthew J. Penn, Neil Scheidwasser, Christl A. Donnelly, David A. Duchêne, Samir Bhatt

## Abstract

Using genetic data to infer evolutionary distances between molecular sequence pairs based on a Markov substitution model is a common procedure in phylogenetics, in particular for selecting a good starting tree to improve upon. Many evolutionary patterns can be accurately modelled using substitution models that are available in closed form, including the popular general time reversible model (GTR) for DNA data. For more unusual biological phenomena such as variations in lineage-specific evolutionary rates over time (heterotachy), more complex approaches uch as the GTR with rate variation (GTR**+Γ**) are required, but do not admit analytical solutions and do not automatically allow for likelihood calculations crucial for Bayesian analysis. In this paper, we derive a hybrid approach between these two methods, incorporating **Γ(*α, α*)**-distributed rate variation and heterotachy into a hierarchical Bayesian GTR-style framework. Our approach is differentiable and amenable to both stochastic gradient descent for optimisation and Hamiltonian Markov chain Monte Carlo for Bayesian inference. We show the utility of our approach by studying hypotheses regarding the origins of the eukaryotic cell within the context of a universal tree of life and find evidence for a two-domain theory.

## 1 Introduction

Phylogenetic distance is a fundamental quantity when considering the evolutionary history of a set of taxa. Its definition serves as the cornerstone for constructing phylogenetic trees, which encapsulate the evolutionary relationships between organisms. These trees, represented as bifurcating binary structures, visually depict the evolutionary distances and genetic relatedness among different species or organisms. In both likelihood and minimum evolution methods, the lengths of the branches - representing the evolutionary time between bifurcations - and the overall tree topology are ultimately heavily dependent on distance calculations, either directly through the objective function or implicitly in the estimation of the model parameters.

Many methods for defining phylogenetic distance exist. The simplest such metric is the Hamming distance which, for a pair of taxa *g* and *h*, simply measures the proportion of sites which take a different value in the two genomes. Although its simplicity is appealing, the Hamming distance ignores the possibility of multiple changes at a given site (“multiple hits”) as well as the heterogeneity in the character set *χ*, which for amino acid or codon sequences is somewhat large.

The solution ubiquitously used to these challenges is to introduce a Markov model for the substitution process, where we suppose that each site evolves according to an independent continuous-time reversible Markov chain (CTMC) with a substitution rate matrix *Q* and a transition matrix *P* (*t*). We suppose that each chain is started at taxon *G* according to the stationary distribution, ***ξ***, and that the corresponding site in genome *H* takes the value of the CTMC at time *d*(*g, h*), the evolutionary distance between this pair of taxa. From this we can use an objective criteria such as minimum evolution (Day et al., 1986; Pauplin, 2000; Desper and Gascuel, 2002; Gascuel and Steel, 2006; Lefort et al., 2015; Penn et al., 2023) to estimate a phylogenetic tree.

For a range of CTMC models, the genetic distance is available in closed form. For example, under the general time reversible model (GTR), the genetic distances are simply

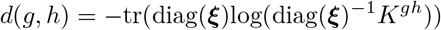

where *K*^*gh*^ is known as the frequency or divergence matrix representing the proportion of changes from the character set *χ*. When incorporating Γ(*α, α*)-distributed rate variation it is possible to define a formula for the GTR+Γmodel (Gu and Li, 1996; Yang and Kumar, 1996) which can be represented in closed form via eigendecomposition, but does not admit an analytical solution. To estimate distances from a GTR+Γmodel, several approaches have been suggested. It is possible to use empirical rates for the GTR substitution matrix via a simple average (Waddell and Steel, 1997; Gatto et al., 2007), but these can be problematic due to negative eigenvalues, which means that heuristic adjustments are needed. For the Γparameter, *α*, parsimony can be used (Yang and Kumar, 1996). More recently, the parameter *α*, distances and GTR parameters can all be considered free parameters and optimised via gradient descent (e.g. Penn et al. (2023)), but the number of distances grows quadratically with the number of taxa, and therefore optimisation can become intractable quickly. Additionally, for protein residues, the larger character set means there is a risk of overfitting. Alongside the GTR model, another popular alternative is the log-det or paralinear model (Lake, 1994; Lockhart et al., 1994), which calculates a genetic distance solely from *K*^*gh*^ and includes a geometric mean estimator for the frequencies that allows the model to account for heterotachy.

Here we build on the work of Lake (1994); Lockhart et al. (1994); Yang and Kumar (1996); Gu and Li (1996); Waddell and Steel (1997); Gatto et al. (2007); Penn et al. (2023), assimilating their previous results to create a new Bayesian model to estimate genetic distances under a GTR+Γ with heterotachy. We first introduce the concept of frequency matrices and rederive the log-det estimator in full. We then rederive an equation for the analog to the log-det estimator under Γ variation using a simple spectral approach first introduced by Gu and Li (1996). This results in a closed-form formula for the distance based only on *α*. In previous research (Gu and Li, 1996), *α* is independently estimated and no provisions for heterotachy are made. We present an approach that can model GTR+Γ with heterotachy and utilises the standard Markov likelihood for distance models to learn *α* and the GTR rates from the data. We then introduce a Bayesian hierarchical model that puts a prior distribution on GTR rate parameters and *α* but uses an estimator for ***ξ***. Through implementing SVD decompositon, we mitigate the numerical issues from using GTR rates computed directly from the data (Gatto et al., 2007), ensuring that we always get a valid eigendecomposition. Moreover, allowing ***ξ*** to vary from pair to pair, our model also accounts for heterotachy. Thus, our model is a hybrid between the log-det-style models and common estimation approaches for the standard GTR+Γ from Gu and Li (1996).

## 2 Methods

### 2.1 Setup

We consider a set of *n* taxa and suppose that each taxon has a (random) genomic sequence *G*^1^, …, *G*^*n*^ with *N* total sites for each of them (though the same methods could easily be applied to, for example, proteomic data).

We suppose that the “baseline” mutation CTMC has Q-matrix *Q* and transition matrix *P* (*t*). When we allow rate variation, we will suppose that the true CTMC at a given site has Q-matrix *τ Q* for some

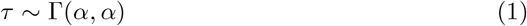

where the value of *τ* is independent of all other sites. Hence, the transition matrix becomes Π(*t*) = *P* (*t/τ*).

Note that the stationary distribution, defined to be ***ξ***, of our CTMC does not depend on the value of *τ* (as if ***ξ***^*T*^ *Q* = 0 then ***ξ***^*T*^ (*Qτ*) = 0 also).

Throughout the majority of this paper, we will work in the large-sites limit. All limiting results for random variables will be under almost-sure convergence.

#### 2.1.1 Frequency matrices

For each pair of taxa (*a, b*), we can define a frequency matrix *K*^*ab*^ such that

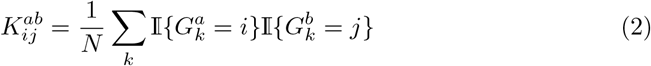

That is, 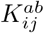 gives the proportion of sites which take value *i* in taxon *a* and take value *j* in taxon *b*. As an example, the sequences

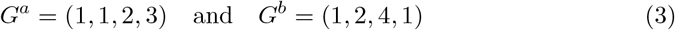

would lead to

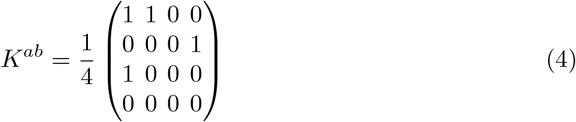

#### 2.1.2 Taking the large-sites limit

Consider the large-sites limit *N* → ∞. As

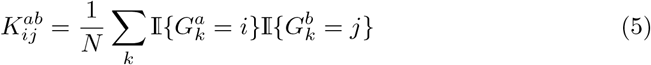

is a sum of independent random variables, we can use the strong law of large numbers (SLLN) to show that

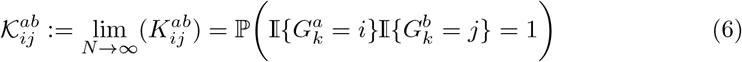

which, if *X*_*t*_ is a copy of the (general, scaled) mutation CTMC and *D*_*ab*_ is the distance between taxa *a* and *b*, can be rewritten as

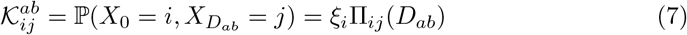

This result relies on the independence of each site. However, extensions of the SLLN exist for various classes of weakly dependent random variables, some of which may be appropriate for phylogenetic modelling (Philipp and Stout, 1975).

For the remainder of this paper, we will assume that *K* approximates 𝒦 well, and will therefore take it to (at least approximately) satisfy (7).

### 2.2 The log-determinant

Firstly, we consider the case with no rate variation, so that Π = *P*. Equation (7) can then be rewritten succinctly in matrix form as

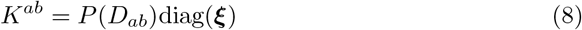

where for a vector ***v***, diag(***v***) refers to a diagonal matrix Δ with non-zero entries Δ_*ii*_ = *v*_*i*_.

We seek to use this equation to find *D*_*ab*_ without relying on the parameters of the CTMC. Note that, by the forward equations for a CTMC,

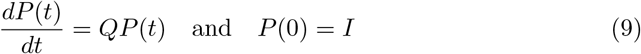

which has a solution

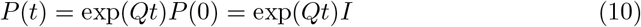

Then, using the fact that for a matrix *A* with eigenvalues *λ*_*i*_

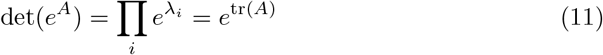

This follows as the eigenvalues of *e*^*A*^ are the exponentials of the eigenvalues of *A*, the determinant is the product of the eigenvalues, and the trace is their sum. Therefore,

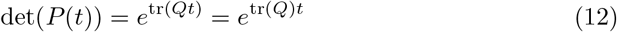

Now, using (8)

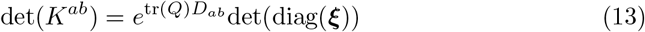

which rearranges to

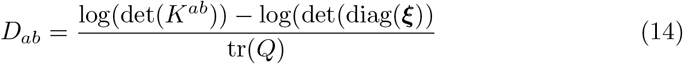

In this model, there is a degree of freedom in the parameters as one can rescale time (which results in multiplying *D* by some 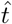 and *Q* by 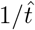). In (14), the term tr(*Q*) can be treated as this scaling parameter, and therefore without loss of generality, we can set it equal to a convenient value. In this paper, we shall choose tr(*Q*) = −*C*, where *C* is the number of possible values that a given site could take (noting that, as it comes from a Q-matrix, tr(*Q*) must be negative). Thus, we have

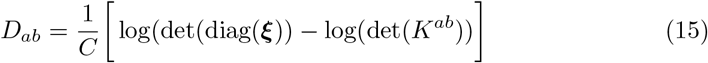

#### 2.2.1 Estimating *ξ*

To complete our independence from the parameters of the CTMC, we must find a way to estimate 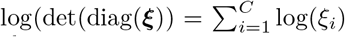. We do this by using the frequency of sites ***F*** ^*a*^ and ***F*** ^*b*^ at each of the taxa where, for example

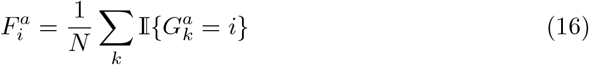

There are a plethora of possible consistent estimators that one could use - perhaps the most natural being the arithmetic mean of frequencies between the pairs *a, b*

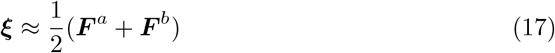

In the log-det formula, we need to estimate log(***ξ***). Thus, it is natural to instead use

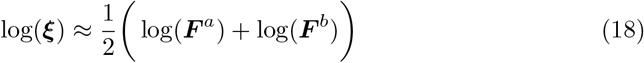

Using this estimator is equivalent to taking the average of the distance estimates resulting from ***ξ*** ≈ ***F*** ^*a*^ and ***ξ*** ≈ ***F*** ^*b*^.

In effect, this relaxes our reversibility assumption, no longer assuming that the distribution of the sites is the same at taxon *a* and taxon *b*. Indeed, one can note that throughout the derivation, we only used the fact that ***ξ*** was the frequency distribution at taxon *a* (although we have tacitly assumed that *Q* is diagonalisable, which is not guaranteed for non-reversible chains). When comparing distant taxa, whose distribution of sites may well not be the same, this can increase the accuracy of distance approximations (Baake, 1998).

#### 2.2.2 The classical log-det metric

Thus, we recover the classical log-det distance, introduced in (Lake, 1994; Lockhart et al., 1994)

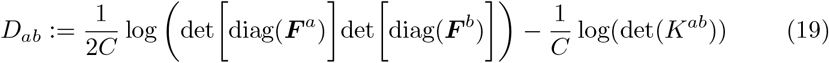

Note that from the almost-sure convergence of *K*^*ab*^ to *𝒦*^*ab*^, the almost-sure convergence of our estimator for ***ξ***, and the continuity of Equation (14), the log-det distance must converge to the true distance in the *N* → ∞ limit.

In the appendix, we explore the metric properties of the log-determinant. We show that for finite *N*, it may not satisfy the triangle inequality, but in the *N* → ∞ limit, it is a true metric.

#### 2.2.3 Rate variation across sites

As previously discussed, it is common to incorporate rate variation across sites. Recall that

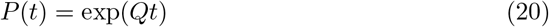

Incorporating rate variation involves integrating over a probability distribution function *P* for the scale *τ*

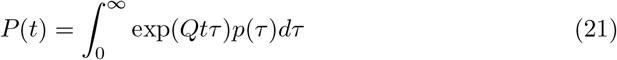

We suppose that *τ* is gamma-distributed with shape and scale *α*. From Lemma 1, we know that *Q* is diagonalisable and so define eigenvalues ***λ*** and eigenvectors *U*. Then,

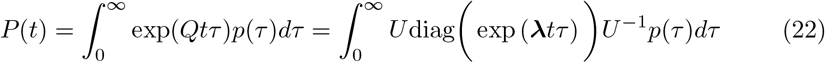

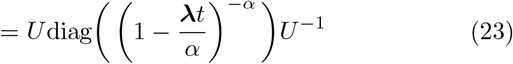

Hence, using the fact that the determinant is the product of the eigenvalues,

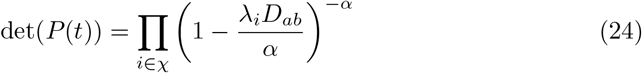

and so

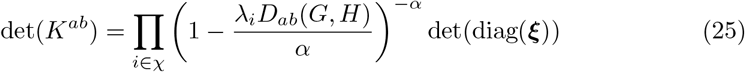

Unlike in the GTR case, it is impossible to infer the distance *D*_*ab*_ without knowing the parameters *λ*_*i*_. Treating these as free parameters in an optimisation scheme does work reasonably well in practice, but inverting these equations carries reasonable computational cost, while accurately finding the *λ*_*i*_ requires many inter-taxa pairs to be used simultaneously.

#### 2.2.4 An alternative approach to estimating distances

However, we can improve on the result in (25) through an eigenvalue approach. Note that

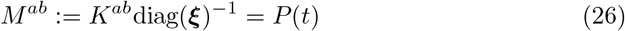

where we can treat *M* as a known matrix by estimating ***ξ*** from the frequency vectors ***F***. We assume that *M* can be diagonalised and has eigenvalues *μ*_1_, …, *μ*_*N*_. Now, equating these to the eigenvalues of *P* (*t*), we get a system of equations

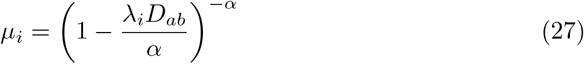

and hence

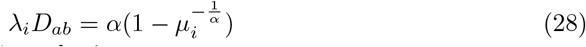

Summing these equations over *i* results in

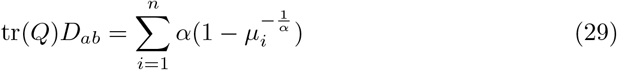

Again setting the trace of *Q* to be equal to *C*, we get

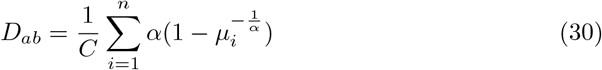

which provides an estimate for the distance that has parametric dependence only on *α*. Note that in the *α* → ∞ limit, we recover

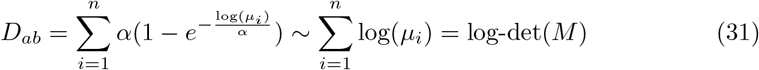

which is the standard log-det metric. This equation was first derived by (Gu and Li, 1996) and has not been used widely. Our novel contribution is to use the idea of (Gu and Li, 1996) to create a non-stationary estimator which can be used within a Bayesian model to allow shrinkage.

#### 2.2.5 Practical eigenvalue calculation

Note that when computing the eigenvalues *μ*_*i*_, *M*^*ab*^ is non-symmetric and can have high condition numbers. To increase stability, we find a symmetric matrix with the same eigenvalues. This can be done by noting that (as shown in Lemma 1), from reversibility, the matrix

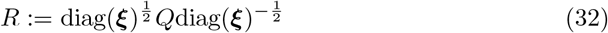

is symmetric. Thus,

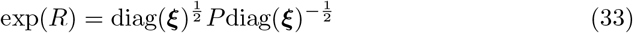

is symmetric, and also similar to *P*, meaning it has the same eigenvalues. We therefore also know that

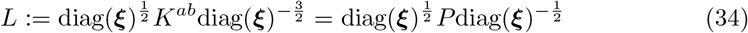

is (at least approximately) symmetric. Thus, we can use this to calculate the values *μ*_*i*_ in a more numerically stable fashion than directly using *M*.

On smaller datasets, even *L* can cause numerical problems due to being poorly conditioned. We have found that it is better to use the singular values of *L* (which, if it were symmetric, would coincide with the eigenvalues) to approximate *μ*_*i*_ in the most stable fashion.

#### 2.2.6 Estimating *ξ*

Again, these distance estimates rely on the value of ***ξ*** and so this must be approximated. As before, we take the average of the distances given by using ***ξ*** ≈ ***F*** ^*a*^ and ***ξ*** ≈ ***F*** ^*b*^. Thus,

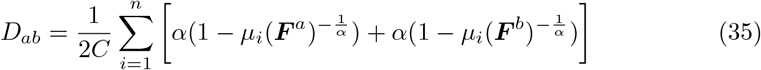

Again, this reduces our reliance on the reversibility of the CTMC (though to use the SVD method discussed in the previous section, reversibility is necessary to guarantee the symmetry of *L*). Practically, particularly on smaller datasets, we have found that using

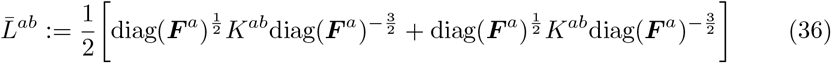

and then setting *μ*_*i*_ to be the eigenvalues of 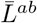 can reduce the numerical instability (as a trivial example, if one of the ***F*** ^*·*^ contains a 0, then we will not be able to calculate an *L*-matrix from that vector). On the datasets considered in this paper, both of these methods give similar results, but we present the results of this second method in our examples.

#### 2.2.7 Bayesian distance estimation

Our approach can be easily nested within a Bayesian hierarchical model where prior probabilities can be placed on *α* and rate matrix *Q*. Given our set of genetic sequences we first calculate

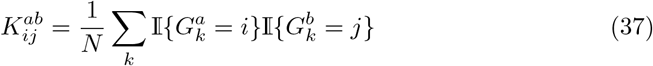

and

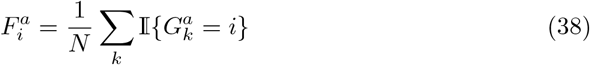

for each pair of taxa *a* and *b*. Note ***F*** ^*a*^ and ***F*** ^*b*^ are simply the row and column sums of *K*^*ab*^.

Using these inputs, which need only be calculated once (and in a manner that can be parallelised), we can then define a hierarchical Bayesian model. Our parameters are assigned prior probabilities as follows

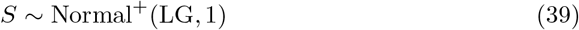

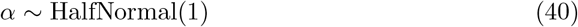

We choose the LG model (Le and Gascuel, 2008) as the mean in our Normal distribution prior over a GTR (Tarvaré, 1986) rate matrix. To make the GTR model symmetric and eigendecomposable we make the substitution matrix *Q* from the parameter rates *S* (which are shared across all pairs), and a pair-specific stationary distribution ***ξ***^*ab*^ given by

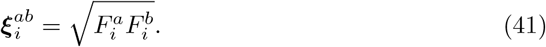

An eigendecomposition on a symmetric form of *Q* therefore yields a unique substitution matrix for each taxon pair, i.e. *Q*^*ab*^ = *U*^*ab*^Λ^*ab*^(*U*^*ab*^)^−1^. From this decomposition, we can compute the probability of a path length between taxa *a* and *b* under Gamma rate variation among sites as 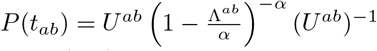. This decomposition needs to be performed for all pairs (*a, b*), and is the main bottleneck in our approach.

Next we compute the matrix

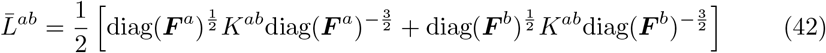

From which we can define the following singular value decomposition *U*^*ab*^Σ^*ab*^*V* ^*ab*^ = 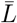 from which the distance

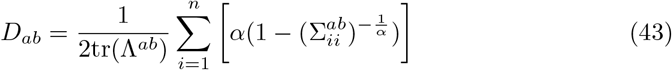

Note, these singular values are computed only once. Finally, our log-likelihood is

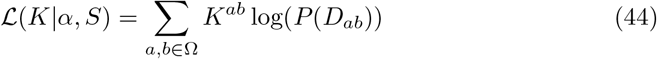

Using this likelihood with our prior probabilities we can define a posterior distribution *p*(*α, S*|*K*) which is differentiable and can therefore use state-of-the-art Markov chain Monte Carlo (MCMC) samplers such as Hamiltonian Monte Carlo (Hoffman and Gelman, 2014). For each posterior sample of distance matrices formed from *d*(*a, b*), a tree can be estimated.

Note that, throughout this Bayesian fitting, we use the assumption that our model is time-reversible. This is necessary both to avoid overfitting (as it reduces the number of parameters) and to reduce the computational cost. However, we still use our reversibility-agnostic estimates for the inter-taxa distances given the parameter *α*. This helps to make our model as flexible as possible, while remaining within the practical constraints of the GTR framework.

## 3 Results and discussion

### 3.1 A simulated example

To begin, we consider an example based on simulated data. We sample trees uniformly at random, with 50 taxa and with uniform branch lengths. We rescaled root-to-tip distances to 0.3, which is a sensible value for real data (Klopfstein et al., 2017). We them simulate amino acid sequences down this tree with 10,000 sites under an LG model, and a 4-category discrete Gamma distribution with shape sampled uniformly between 0.1 and 5l. We optimise the log-likelihood in Equation (44) using L-BFGS-B optimisation. In addition, we also treat frequencies and rates has free parameters and optimise a full GTR model. This full GTR model has a free parameter for every pair of taxa and therefore quickly becomes unfeasible for large trees. Figure 1 shows that our approach and the GTR model results in virtually the same likelihood as for the true data and the resultant tree are close to the ground truth for both models.

**Fig. 1.**
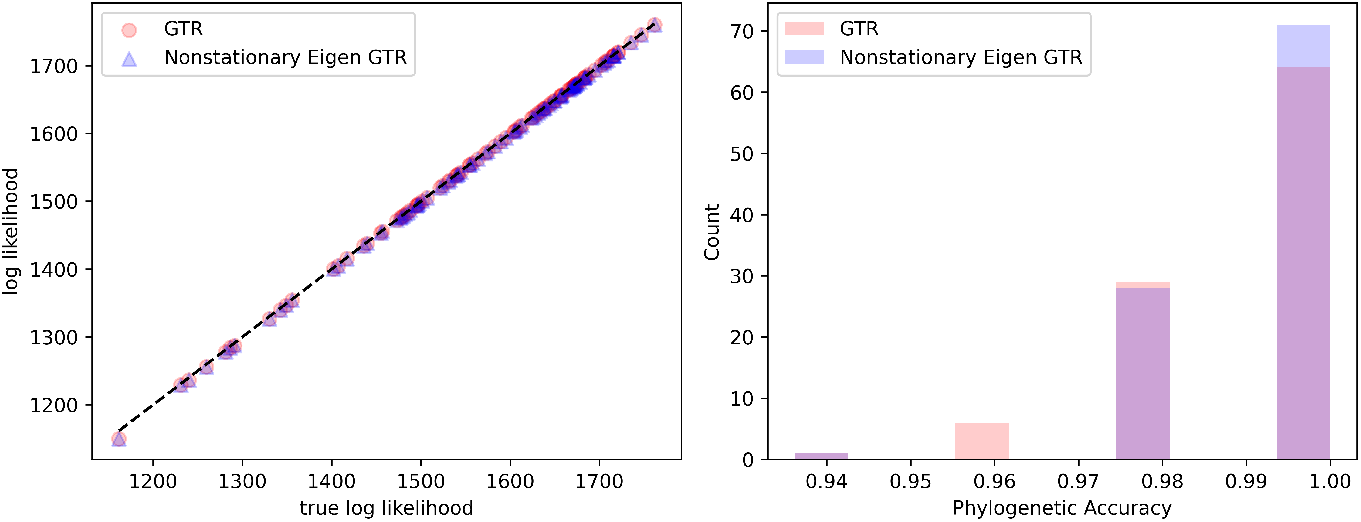
(left) Comparison of our proposed non-stationary Eigen GTR and a full GTR trained with free parameters for every taxa pair with the true log-likelihood for the actual data. (right) Phylogenetic accuracy measured as one minus the RF distance to the true tree from both the non-stationary Eigen GTR and full GTR

### 3.2 Applications to a protein-based dataset

Secondly, we apply our approach to the dataset and problem introduced by Williams et al. (2019) studying hypotheses regarding the origins of the eukaryotic cell within the context of a universal tree of life. Using their dataset comprised of 35 core genes (∼ 7000 protein residues) across 83 taxa, we can apply our Bayesian eigenvalue approach. In their original analysis, Williams et al. (2019) find a GTR + Γ+ F model yields strong support for a three domain tree of life, but discuss idiosyncrasies in the data that necessitate more sophisticated models of molecular evolution, and under these find stronger statistical support for a two-domain tree of life. Analysing the same data as Williams et al. (2019) we first note that a basic log-det distance estimator (Lake, 1994) results in several negative branches between taxa and therefore is invalid. Using an LG model and recovering a bootstrapped distance-based tree via balanced minimum evolution in FastME (Lefort et al., 2015), we find that this tree strongly supports a three-domain tree of life. However, applying our new Bayesian distance-based algorithm, using a GTR model but with an LG prior, results in a two-domain tree (see Figure 2) with 100% posterior support. While differences exist between our tree and the tree from Williams et al. (2019), and in general we do not expect distance-based approaches to match those based on Felsenstein’s likelihood, our approach shows that distance-based approaches can include important molecular evolution processes such as rate variation and heterotachy. Importantly, due to the ability to utilise state-of-the-art MCMC approaches, our posterior which included 2000 samples, and two chains shows excellent convergence with no pathologies (see Figure 3).

**Fig. 2.**
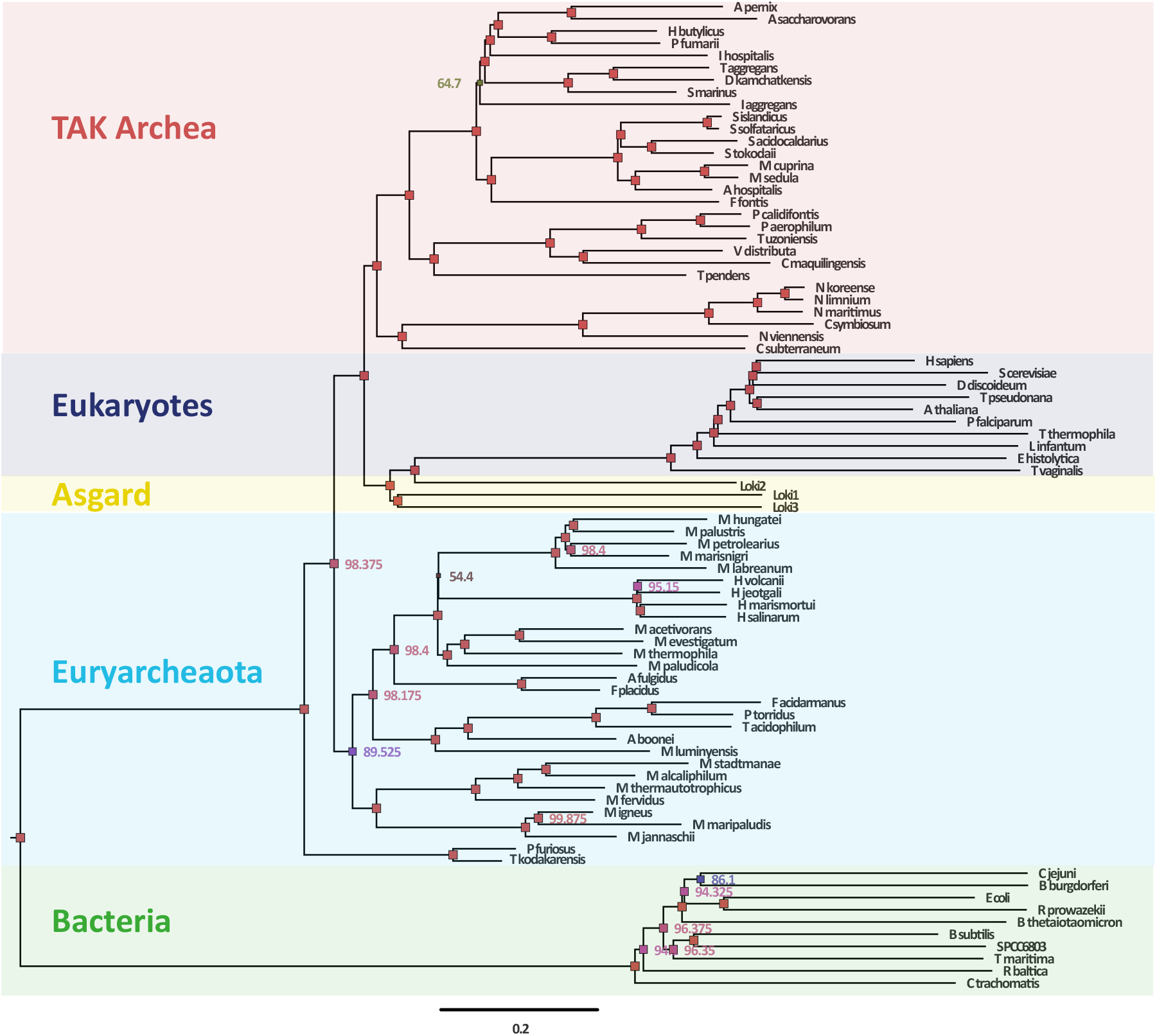
Reanalysis of Williams et al. (2019) studying hypotheses regarding the origins of the eukaryotic cell within the context of a universal tree of life. Using our GTR+Γwith hereotachy and prior shrinkage, we find support for two domains of life

**Fig. 3.**
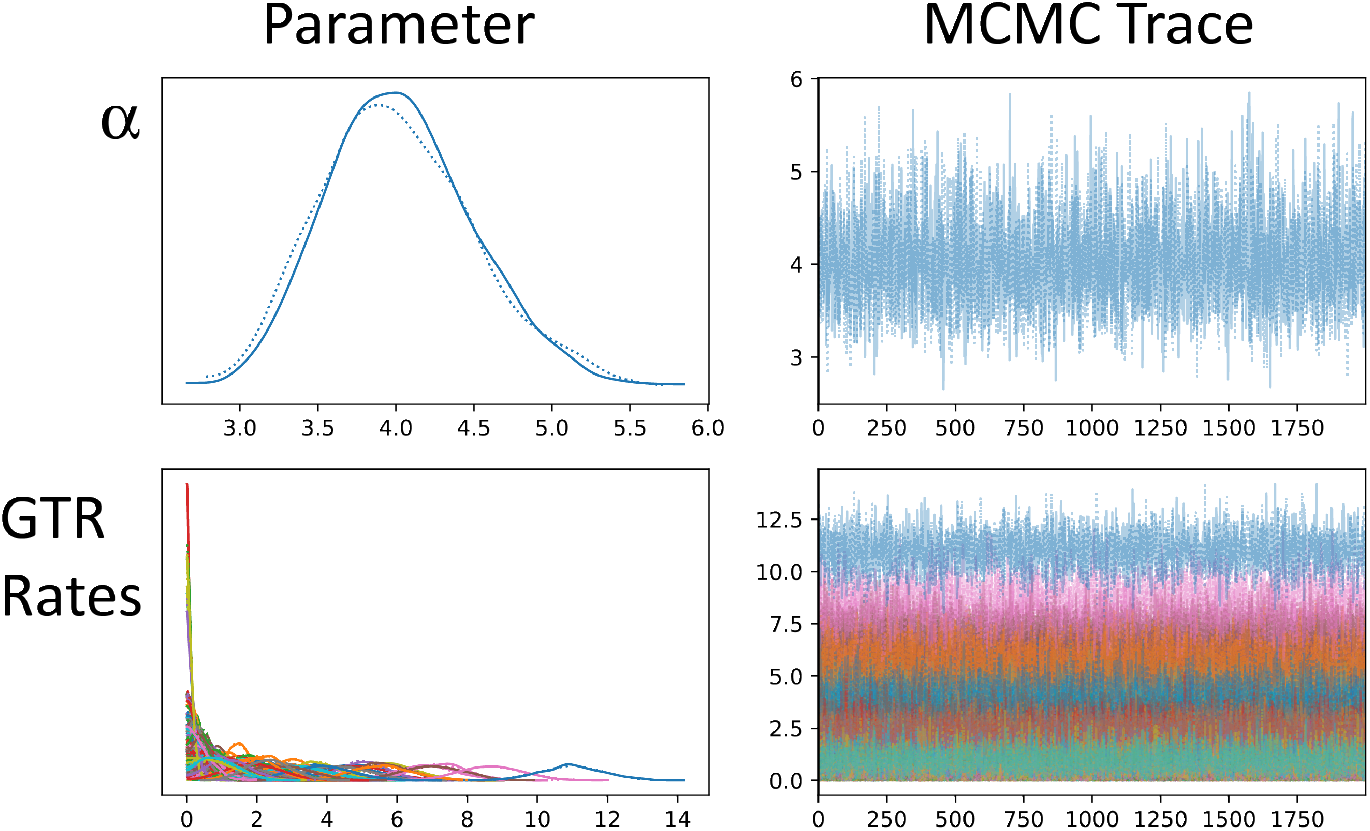
MCMC trace file showing excellent convergence.

### 3.3 Conclusions

The method introduced in this paper provides a flexible and reliable way to estimate the phylogenetic distances between a set of taxa. By combining ideas from the GTR and log-det approaches, our model has the flexibility to incorporate heterotachy across the dataset and calculate a likelihood for each set of parameters while also being practically computable. Particularly when looking at large-timescale datasets, these properties are vital in ensuring that our phylogenetic analysis is accurate, both in terms of the central estimate and the uncertainty around it.

We hope to consider extensions to this model in future work, potentially considering different distributions that could more accurately, or more flexibly, describe the variation in genetic rates. Furthermore, given the wide range of possible estimators for the stationary distribution ***ξ***, we hope to investigate the different possibilities more systematically in order to perhaps improve the version used in this paper.

## 4 Competing Interests

The authors declare no competing interests.

## 5 Funding

M.J.P acknowledges support from his EPSRC DTP studentship, awarded by the University of Oxford to fund his DPhil in Statistics. S.B. acknowledges funding from the MRC Centre for Global Infectious Disease Analysis (reference MR/X020258/1), funded by the UK Medical Research Council (MRC). This UK funded award is carried out in the frame of the Global Health EDCTP3 Joint Undertaking. S.B. is funded by the National Institute for Health and Care Research (NIHR) Health Protection Research Unit in Modelling and Health Economics, a partnership between UK Health Security Agency, Imperial College London and LSHTM (grant code NIHR200908). Disclaimer: “The views expressed are those of the author(s) and not necessarily those of the NIHR, UK Health Security Agency or the Department of Health and Social Care.” S.B. acknowledges support from the Novo Nordisk Foundation via The Novo Nordisk Young Investigator Award (NNF20OC0059309). S.B. acknowledges support from the Danish National Research Foundation via a chair grant (DNRF160) which also supports N.S. S.B. acknowledges support from The Eric and Wendy Schmidt Fund For Strategic Innovation via the Schmidt Polymath Award (G-22-63345). S.B and N.S acknowledge the Pioneer Centre for AI, DNRF grant number P1 as affiliate researchers. C.A.D receives support from the NIHR HPRU in Emerging and Zoonotic Infections, a partnership between the UK Health Security Agency, University of Liverpool, University of Oxford and Liverpool School of Tropical Medicine (grant code NIHR200907).

D.A.D. is funded by a European Research Council Marie Sklodowska-Curie fellowship (H2020-MSCA-IF-2019-883832).

## Appendix A Diagonalisability of *Q*

### Lemma 1.

*The matrix Q is diagonalisable*

**Proof:** Note that, as *Q* is reversible

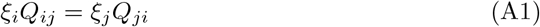

Hence, this means that the matrix 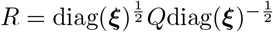 is symmetric as its entries are

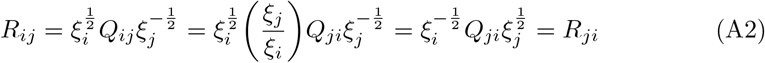

Thus, *R* is diagonalisable and so there exists a diagonal matrix *D* and an invertible matrix *V* such that

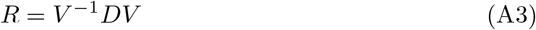

and hence

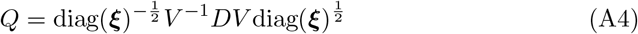

and hence, setting 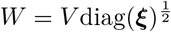, we see that

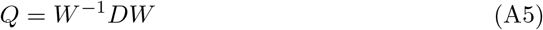

as required for diagonalisability.

## Appendix B Metric properties of the log-determinant

A metric *D* on a set *S* must satisfy three properties

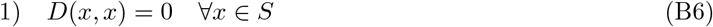

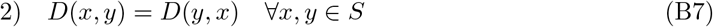

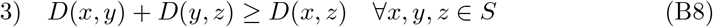

The log-determinant distance *D*_*ab*_ always satisfies the first two of these properties, with 1) following as, when *a* = *b*, we have

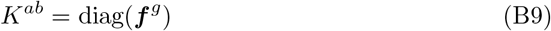

and so we see *D*_*aa*_ = 0.

The second property also follows quickly as swapping *a* and *b* simply transposes *K*^*ab*^, therefore leaving its determinant unchanged.

However, for finite *N, D* may not satisfy 3). For example, it is undefined if the determinant of *K* is non-positive (as a trivial case, this always happens when *N <* 4 as one row will only contain 0’s). Even when restricted to positive determinants, the triangle inequality can fail - for example, with the sequences

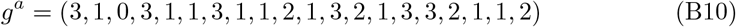

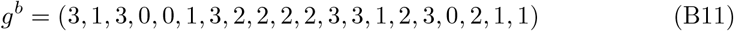

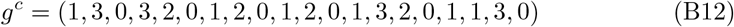

which gives

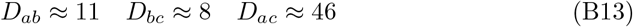

and hence

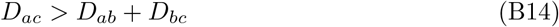

Thus, care must be taken when applying this metric to small phylogenetic datasets, although, due to the generally small state-space *χ* compared to large *N*, this distance will give good performance on phylogenetic datasets.

Note that in the the *N* → ∞ limit, *D* does satisfy 3) if evolutionary distance also satisfies 3). In a general phylogenetic tree construction, this holds - given three points, *A, B* and *C* on a tree (representing taxa - though they need not be nodes here), their evolutionary distance is simply the distance between them on the tree, and therefore it is simple to show that *D* is a metric. Briefly, one can use the fact that there is a unique path between any pair of points. The path between *A* and *C*, 𝒫 (*A, C*) can be found by choosing all points that are on exactly one of 𝒫 (*A, B*) and 𝒫 (*B, C*). Thus, the length of 𝒫 (*A, C*) is necessarily shorter than the sum of the lengths of 𝒫 (*A, B*) and 𝒫 (*B, C*) as required.

## Notes

### Competing Interest Statement

The authors have declared no competing interest.

